# Delayed copulation and mating in the malaria vector *Anopheles funestus* compared to *Anopheles arabiensis*

**DOI:** 10.1101/2025.01.20.633855

**Authors:** Emmanuel Elirehema Hape, Alex Thadei Ngonyani, Daniel Mathias Mabula, Joel Daniel Nkya, Claus Augustino Thomas, Mohamed Jumanne Omari, Doreen Josen Siria, Halfan Said Ngowo, Lizette Leonie Koekemoer, Fredros Oketch Okumu

## Abstract

Mating is a vital behavior for mosquito reproduction, yet it remains poorly understood under captive conditions. We examined the copulation dynamics of two key malaria vectors, *Anopheles funestus*, and *Anopheles arabiensis*, in controlled laboratory settings in Tanzania. We observed how variations in mosquito age and artificial lighting influence mating success for these two mosquito species within cages under controlled conditions. We conducted observations in 24-hour cycles, monitoring copulation events and insemination in females. We used generalized linear mixed models (GLMMs) for statistical analyses to assess how environmental conditions influence mating behavior. We found that *Anopheles arabiensis* exhibited rapid copulation, with 32.4% of individuals mating by Day 3 post-emergence, while *An. funestus* showed delayed activity, reaching a similar mating rate by Day 8. The introduction of artificial red light significantly accelerated copulation in *An. funestus* but did not affect *An. arabiensis*. Dissection confirmed successful sperm transfer and mating plug delivery in over 92% of copulating pairs for both species. Mating occurred primarily at night, with distinct peaks at 22:00 for *An. arabiensis* and 23:00 for *An. funestus*. In conclusion, our findings reveal species-specific differences in reproductive behavior, which could improve the colonization of *An. funestus*, a species historically challenging to rear in captivity. These insights may also inform the development of new vector control technologies, such as sterile insect techniques and genetic-based approaches, that exploit mosquito mating behaviors.

## Background

Vector control methods, particularly through insecticide-treated nets (ITNs) and indoor residual spraying (IRS), remain the cornerstone of malaria control in Africa but are increasingly challenged by adaptive mosquito behaviors and widespread insecticide resistance (Bhatt et al., 2015; Sanou et al., 2021; Odero et al., 2024). While much is known about the biting and feeding patterns of *Anopheles* mosquitoes, critical gaps remain in understanding their copulation and mating behaviors, particularly those of key vectors such as *Anopheles funestus* and *An. arabiensis* (Carrasco et al., 2019; Takken et al., 2024).

Mating behavior in *Anopheles* mosquitoes in the wild typically involve swarming, where males aggregate and females enter these swarms to mate (McIver, 1980; Gary et al., 2009; Sawadogo et al., 2014; Kaindoa et al., 2017; Kaindoa et al., 2019; Charlwood, 2023), and is influenced by environmental factors such as light, temperature, and humidity (McIver, 1980; Howell and Knols, 2009). However, the specifics of these behaviors and their variation among species are poorly understood, particularly under natural and semi-natural conditions (Paton et al., 2013; Facchinelli et al., 2015). For instance, *An. funestus*, a highly efficient malaria vector, presents significant challenges for laboratory studies due to difficulties in establishing colonies, with only a few strains, such as FUMOZ from Mozambique in 2000, FANG from Angola in 2002 (Hunt et al., 2005), and recently FUTAZ from Tanzania, being successfully maintained for extended generations since 2020 (Ngowo et al., 2021). In contrast, species like *An. gambiae* and *An. arabiensis* are more easily colonized, providing greater opportunities for detailed behavioral studies (Coetzee and Fontenille, 2004; Nepomichene et al., 2017; Ngowo et al., 2021; Kahamba et al., 2022).

Understanding copulation dynamics is essential for developing innovative vector control strategies, such as the sterile insect technique (SIT) and gene drives (Benelli and Beier, 2017). These methods rely heavily on manipulating reproductive success, with SIT focusing on releasing sterile males to compete with wild males and gene drives aiming to propagate specific genetic traits that suppress populations or block disease transmission (Helinski et al., 2008; Dame et al., 2009). The success of such approaches depends on understanding factors that influence mating success, including male competitiveness, female choice, and the impact of environmental cues (Mclver, 1980; Howell and Knols, 2009). In areas where multiple *Anopheles* species coexist, interspecific mating and potential hybridization could further complicate control efforts, particularly if genes conferring insecticide resistance or other adaptive traits are transferred (Slotman et al., 2004; Choochote et al., 2014; Niang et al., 2015).

Behavioral flexibility among *Anopheles* species also contributes to residual malaria transmission. For example, *An. arabiensis* frequently rests outdoors and exhibits opportunistic feeding on humans and animals, evading traditional indoor-based interventions like ITNs and IRS (Carrasco et al., 2019). These behaviors, coupled with the species’ mating strategies, necessitate a broader focus on their reproductive ecology. Similarly, *An. coluzzii*, which thrives in urban environments, demonstrates ecological adaptability that supports its role as a significant malaria vector in densely populated regions (Niang et al., 2015; Kamau et al., 2024).

Studies under controlled conditions have begun to shed light on the mating systems of these vectors, revealing the importance of population density, sex ratios, and environmental factors in determining mating outcomes (Facchinelli et al., 2015). Such research highlights the importance of targeting reproductive behaviors to enhance vector control. For example, SIT programs must ensure that sterile males effectively compete for mates, while gene drives require a comprehensive understanding of mating networks to achieve efficient dissemination of modified traits (Dame et al., 2009). Moreover, exploring factors such as light intensity and wavelength, which influence swarming and copulation activities, could inform the design of interventions that disrupt mating success in natural settings (Rowland, 1989; Gibson, 1995; Sheppard et al., 2017; Hellhammer et al., 2022; Finda et al., 2023).

Given the challenges posed by insecticide resistance and behavioral adaptations, integrating reproductive behavior research into malaria control programs offers a promising avenue for reducing mosquito populations and disrupting transmission. By advancing our understanding of *Anopheles* mating dynamics, particularly for less-studied species like *An. funestus* and *An. arabiensis*, we can develop more targeted and sustainable strategies to combat malaria in regions like Tanzania and beyond. In this study, we therefore investigated the copulation dynamics of *An. funestus* and *An. arabiensis* in Ifakara, Tanzania, with a primary focus on how mosquito age and light intensity influence mating success.

## Material and Methods

### Mosquitoes

The experiments were conducted using (a) laboratory-colonized *An. arabiensis* maintained since 2009 (Batista et al., 2019), (b) laboratory-colonized *An. funestus* from Tanzania (FUTAZ) maintained since 2020, and (c) laboratory-colonized *An. funestus* from Mozambique (FUMOZ), originally colonized in South Africa and maintained at Ifakara since 2018.

### Laboratory observations of the effects of mosquito age on copulation events in An.funestus and An. arabiensis in captivity

We first investigated the copulation dynamics in *An. arabiensis* and in both FUMOZ and FUTAZ strains of *An. funestus* mosquitoes. This study involved 24-hour observations, with three replicates per species and strain, inside 30 × 30cm cages containing approximately 1500 mosquitoes in sex ratio of 1 female for every 2 males. Observations were made continuously for 16 consecutive days by counting the copular observed, starting the first day post-emergence until copulation activities ceased. The first set of experimental observations was done following the 12h:12h photoperiod whereby a day-time period was initiated by turning on the light (36W, AC220-240V, 50/60Hz) at 07:00 hours, and turning them off during the night starting 18:00 hours. This room and set-up did not allow for gradual sunset and sunrise conditions. In this first set of experiments, the copulation activities during the night-time were observed and counted by using a hand-held flashlight (1W high power LED; HL-558).

In the second experiment, we aimed to determine whether the mating observations were influenced by the flashlight used in previous experiments. To address this, a 7W red light bulb was used instead, following the same photoperiod cycle but with the red light turned on at 18:00 hours during the observation periods. Observations were also continued during the daytime from 06:00 hours when the red light turned off. The red light was chosen because it closely mimics the wavelengths present during sunset, which are less likely to disrupt mosquito behavior (Sheppard et al., 2017; Hellhammer et al., 2022). No additional flashlight was used to observe copulation activities under these red-light conditions.

A HOBO Temp/RH/Light/Ext-Analog Data Logger^@^ was used to record light intensity (LUX) and capture continuous light measurements. The natural light environment in the field was measured as a control, by placing a similar HOBO meter under the tree. However, only light levels were recorded without observing swarming activities, previously reported to occur around 6:40 PM for approximately 12 (±5 SD) minutes (Kaindoa et al., 2017; Kaindoa et al., 2019).

### Laboratory confirmation the successful insemination and transfer of mating plugs to the females captured in copula, and of the sexual maturity of the copulating males

To confirm the mating success, successful insemination and transfer mating plugs, and male sexual maturity, a random number of intact copulas were retrieved quickly from the cage during the observations and killed by freezing for 10 minutes (Figure 1). Observations were conducted without the red light, and hand-held flashlights were used to assist the procedure. The mating success was confirmed by dissecting the females to assess insemination. To do this, the terminalia and last abdominal segment (segment IX) were cut open in saline to expose the spermatheca capsule. Slide mounts of spermatheca capsules were inspected using a compound microscope at 10× magnification for the presence of sperm (Ngowo et al., 2021). A stereo microscope was used to assess the transfer of the mating plug and the completion of the 180^0^-degree rotation of males in the copula (Oliva et al., 2011).

**Figure 1:**
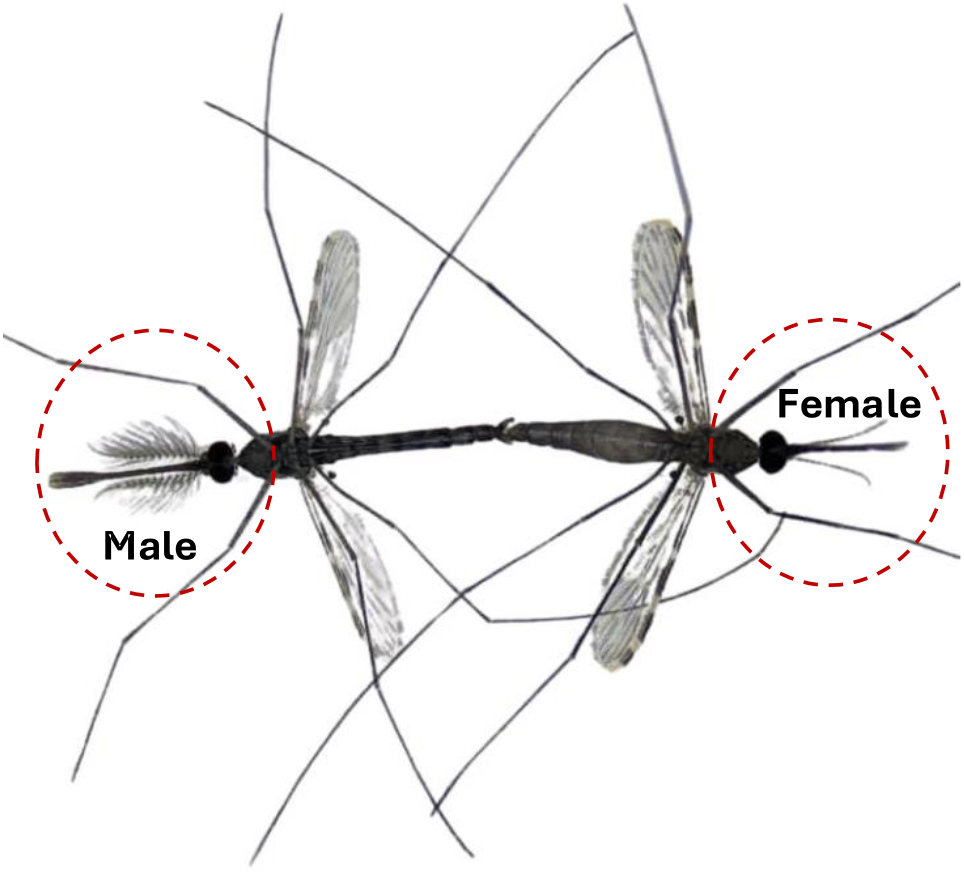
Male-Female Copulation: A male and female *Anopheles* mosquito in copula, demonstrating the typical mating posture where the male clasps the female during swarming behavior.

### Laboratory analysis of peak mating times in mosquitoes at optimal mating age

Adult mosquitoes reared under controlled laboratory conditions were observed continuously to study their peak mating activities over a 24-hour cycle. Mosquitoes were first reared to the age identified as optimal for maximum copulation, based on findings from the observations described above. Three replicates of the observations were conducted for each mosquito species and strain. Swarming and copulation activities in captivity were monitored continuously, starting at 6:00 PM and extending through the night and following day, completing a full 24-hour observation period.

### Statistical analysis

Data collected were statistically analysed using R-software Version 4.4.1; to investigate the impact of different parameters of interest, such as mosquito age (Days), light type, and light intensity on the copulation of different *Anopheles* strains (*An. arabiensis*, FUTAZ, and FUMOZ). Generalized Linear Mixed Models (GLMMs), using the R package, “*lme4”* (Douglas Bates et al., 2024) were used to estimate the mosquito age (Days) at which the maximum copulation activities occurred, as well as the effects of light-type on copulation. The number of copulas observed was modelled following a “*Poisson*” distribution with mosquito strain and light-type as the fixed effect of interest. An experimental replicate was included as the random effect in each of the models generated. Additionally, the proportions of female mosquitoes inseminated, confirmed mating plugs, and males with completed 180^0^ degrees of genitalia rotation retrieved in copula were represented using mean percentages. The peak time of copulation during the 24 hours of observation was calculated using a mean number of copulas. The 24-hour light intensity (LUX) was calculated using the 16-day mean average of the readings obtained using the HOBO Data Logger^@^ in the lab and the field environment. Model selection was done by progressively deleting terms from the maximal model with the “*drop1()*” function, and likelihood ratio tests (LRTs) were employed to assess the relevance of explanatory variables. The “*ggplot2*”, R programmes were used to create all the graphics (Wickham and Chang, 2014). **Results**

### Peak age for copulation and mating

For all *Anopheles* strains, the mean number of copulation events showed a curvilinear relationship with mosquito age (days since emergence). Under conditions without supplementary night light (Figure 2**a**), *An. arabiensis* reached its copulation peak by day 3 post-emergence, accounting for 32.4% of observed copulations. By contrast, *An. funestus* colonies experienced significant delays, with FUTAZ peaking at day 8 (26.3% of copulations) and FUMOZ at day 10 (37.7% of copulations).

**Figure 2:**
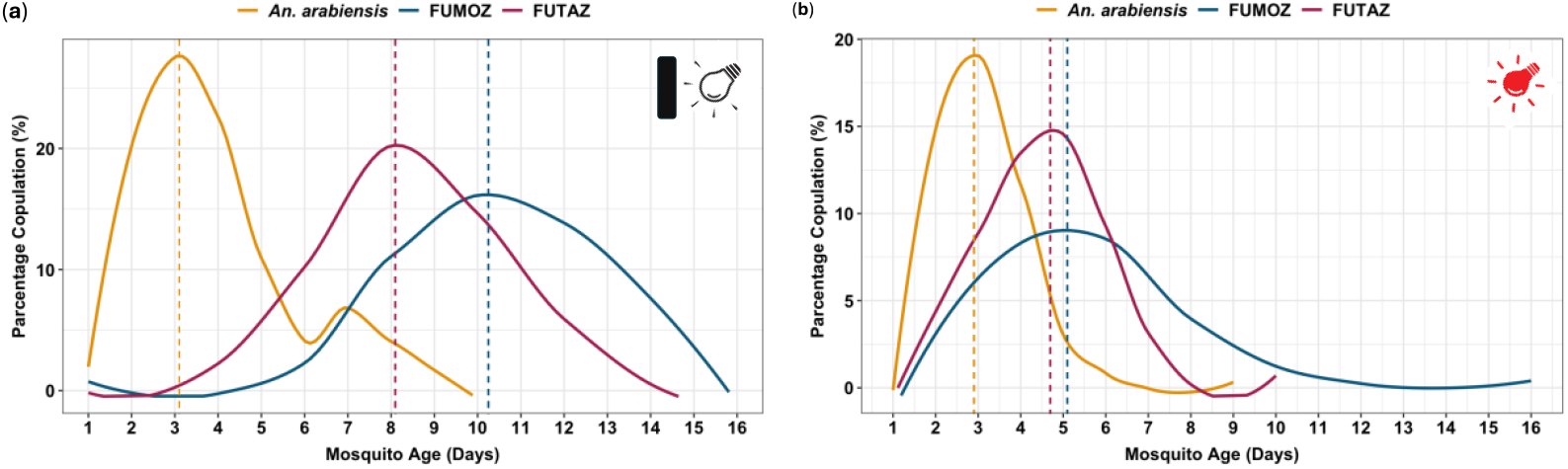
Observation of male-female copulation events in caged mosquitoes. (**a**) Copulation events recorded under dark conditions using a flashlight for counting. (**b**) Copulation events recorded under a 7-watt red light, enabling observation and counting without the need for a flashlight. The vertical dashed lines indicate the optimal age for maximum copulation activity for the specific species and strains.

The use of red light in the insectary did not have a significant impact on the copulation dynamics of *Anopheles arabiensis*. However, light type (χ^2^ =7582.2, df=4, *p*<0.001) significantly reduced the age at which maximum copulation activity was achieved (χ^2^ =4952.3, df=3, *p*<0.001) by both FUMOZ and FUTAZ strains, which reached peak copulation activity approximately 5 days post-emergence compared to conditions without red light (strains: χ^2^ =853.04, df=2, *p*<0.001; Figure 2**b**). In rare observations, male-male copulation was noted in *An. funestus* (Figure 3).

**Figure 3:**
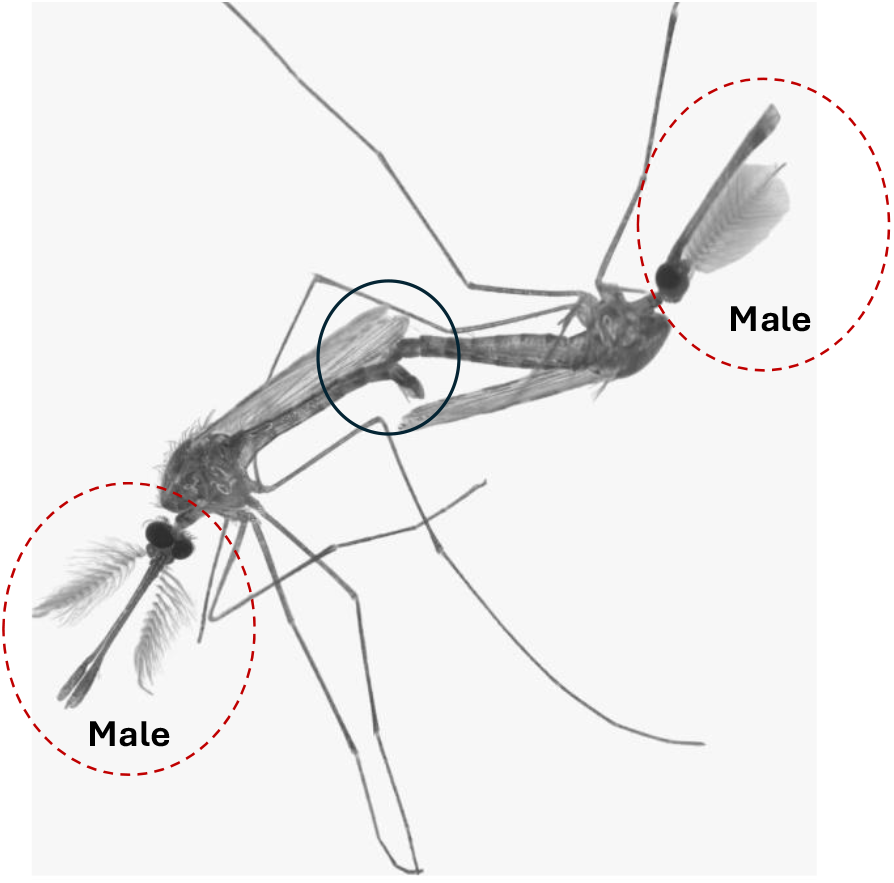
Male-Male Copulation: A rare occurrence, two male *Anopheles* mosquitoes are seen in copula. Unlike typical copulation, one male grabbed the abdomen of the other male rather than the last abdominal segment (black circle). The behavior often resulting from mistaken identity or high competition, highlights the complexity of mosquito mating dynamics under laboratory conditions.

### Observations of successful insemination of females and sexual maturity of the males

More than 92% of dissected female mosquitoes, originally caught in copula, had evidence of successful transfer of sperm to the female spermathecae, along with effective delivery of mating plugs. Moreover, 96 % of the male mosquitoes had completed the 180^0-^degree rotation of their genitalia by the time the copula were retrieved, indicating sexual maturity (Figure 3; Table 1). The number of copula events directly correlated with successful mating events (as indicated by the transfer of mating plugs), and both measures exhibited a curvilinear relationship with mosquito age post-emergence (Figure 4).

**Table 1:** Number of intact copulas collected during the experiment highlighting variability in reproductive outcomes across species.

|  | No. of<br>Copula<br>collected | % Females<br>Inseminated | % Females<br>with<br>Mating<br>Plugs | % Males<br>Sexually<br>Matured |
| --- | --- | --- | --- | --- |
| *An. arabiensis* | 105 | 98.95% | 97.14% | 100.00% |
| *An. funestus* (FUMOZ strain) | 338 | 93.79% | 92.01% | 96.75% |
| *An. funestus* (FUTAZ strain) | 93 | 97.85% | 96.77% | 98.92% |

**Figure 4:**
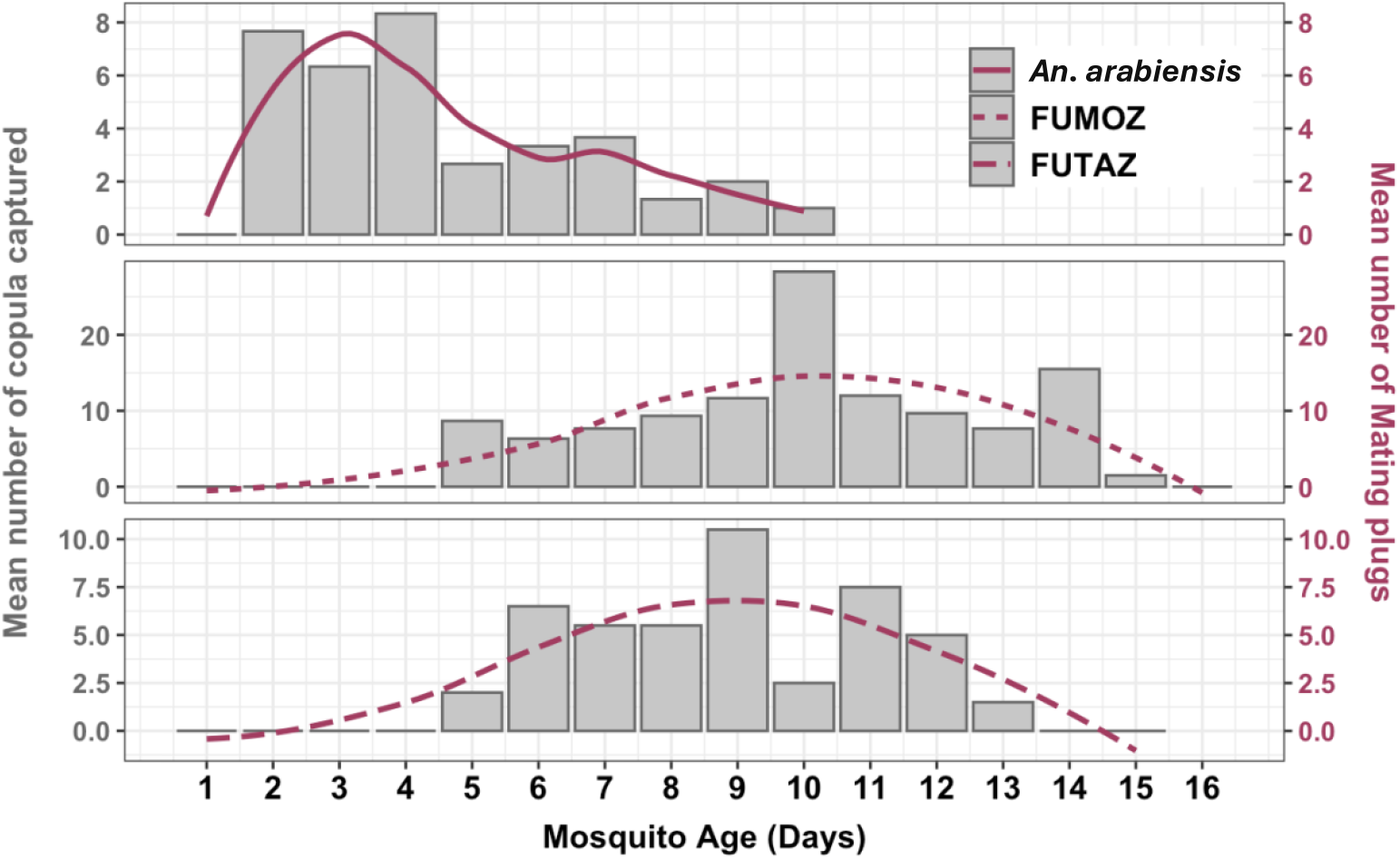
Mating dynamics in *An. arabiensis* and *An. funestus* (FUMOZ and FUTAZ). Over 92% of the randomly collected copulas from different age groups successfully resulted in mating plugs. Observations were conducted using flashlight during total darkness.

### Peak time of copulation and mating

Swarming and copulation activities among mating mosquitoes persist throughout the night though there were differences in peak timing for the species and strains tested. For *An. arabiensis*, the observations showed mating initiation as early as 5 pm, with the peak mating activity sustained until around 6 am, occasionally extending to 7 am. The FUTAZ strain of *An. funestus*, however, demonstrated early mating initiation at 6 pm with more erratic activity patterns throughout the night, but also persisting until 6 am the following morning (Figure-5). In contrast, *An. funestus* (FUMOZ strain) showed a later onset of mating, commencing around 8 pm and reaching peak activity by 11 pm, before concluding by 5 am.

**Figure 5:**
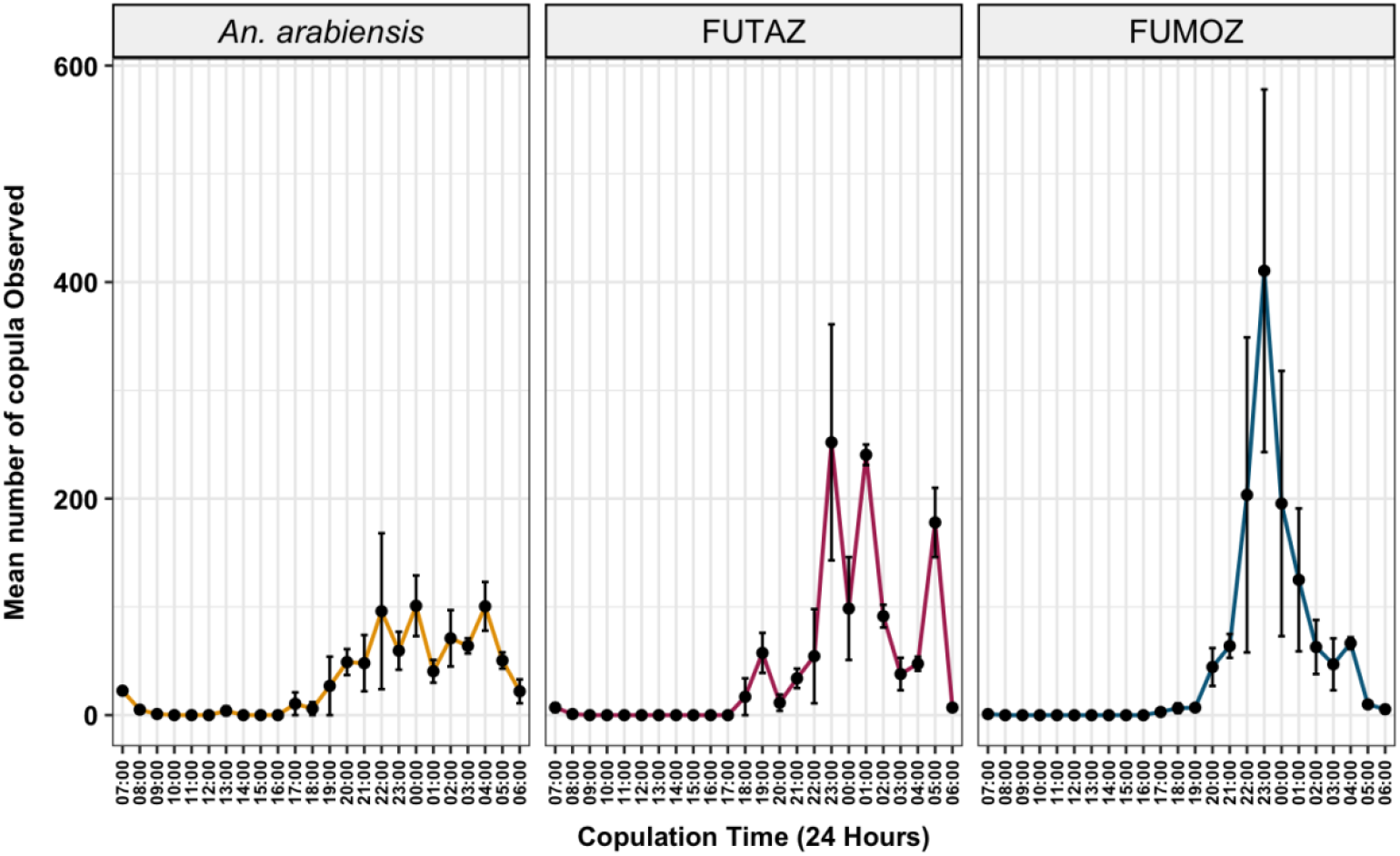
Peak time of copulation and mating, time of day at which peak happens once the mosquitoes attain the peak age of mating.

### Light intensity under copulation events

Under controlled laboratory conditions, light intensity rose from near zero lux at sunrise to roughly 80 lux between 8:00 AM and 5:00 PM, then dropped back to the low levels after sunset unless red light was used, in which case it remained at around 25 lux overnight (Figure-4). In the uncontrolled (field) environment, light intensity began near zero lux at sunrise, exceeded 5,000 lux by midday, peaking around 55000 lux, and then fell below 500 lux around dusk (6:00–7:00 PM), ultimately reaching near zero after 7:00 PM until sunrise (Figure-6).

**Figure 6:**
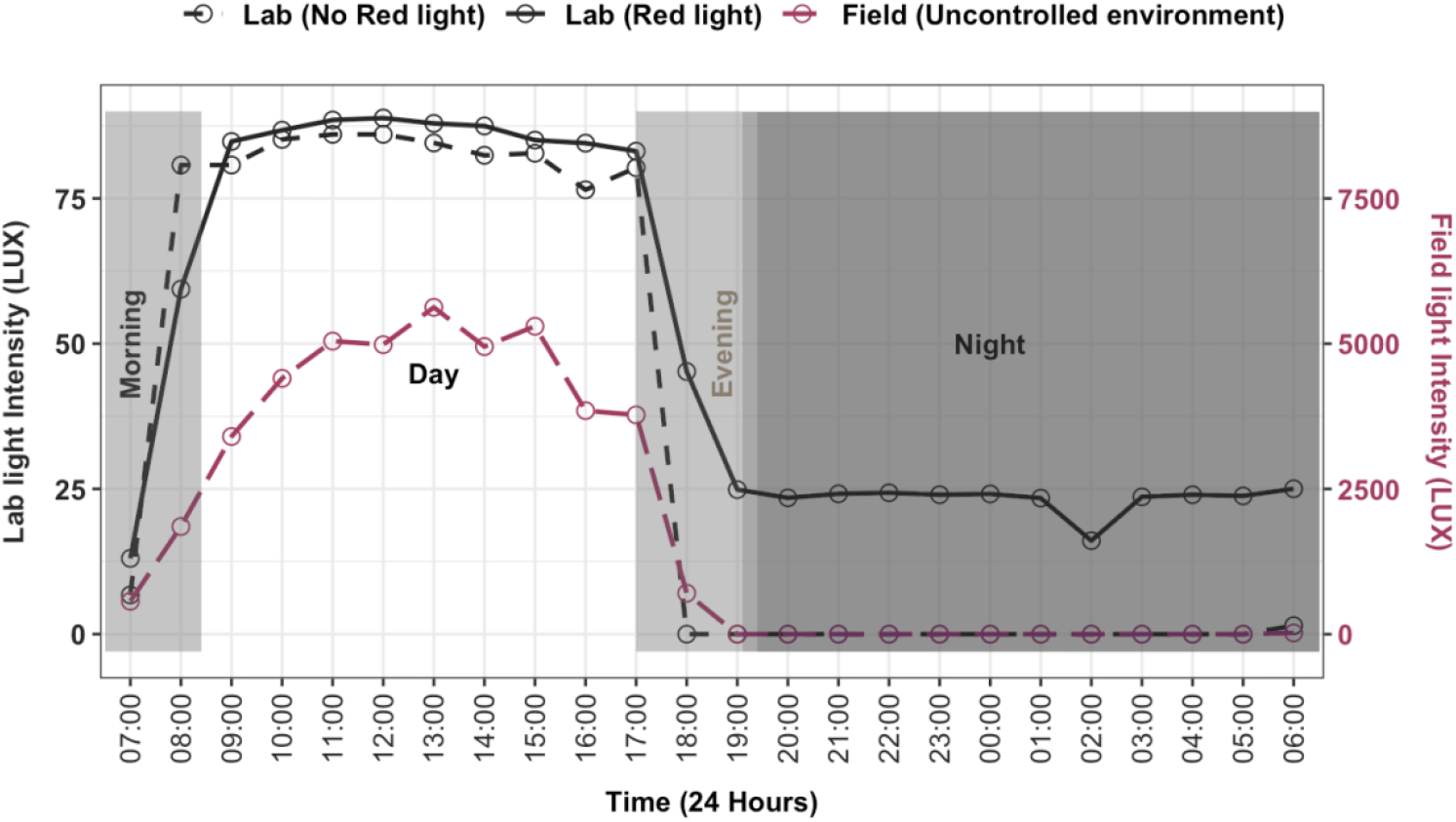
Light intensity recordings: In the lab, gradual transitions between ‘daylight’ and ‘dusk’ phases during light intensity recordings for copulation and mating were not feasible. In contrast, field conditions showed a natural, gradual change.

## Discussion

Understanding the reproductive behaviors of malaria vectors like *An. arabiensis* and *An. funestus* is essential for tracking the population dynamics and developing effective vector control strategies. Mating behaviors, including swarming and copulation, are central to mosquito reproduction, yet they remain understudied, particularly for species like *An. funestus* that are challenging to colonize (Dia et al., 2013; Nepomichene et al., 2017; Kahamba et al., 2022). Swarming and copulation behaviors, influenced by environmental cues such as light intensity, temperature, and humidity, play a pivotal role in mosquito reproduction (Charlwood et al., 2003; Sawadogo et al., 2014; Kaindoa et al., 2019; Baeshen, 2022). This study provides a detailed comparative analysis of copulation dynamics in two key malaria vectors, *Anopheles funestus* and *Anopheles arabiensis*, examining copulation events, insemination, mating plugs, genitalia rotation scores of retrieved copulas, and the effects of field and laboratory light intensity. The observations offer new insights into their reproductive behaviors of these vector species.

Male mosquitoes form swarms at specific times and locations, guided by environmental cues like light, temperature, and humidity. These swarms attract females for mating, with competition and choice determining reproductive success (Charlwood et al., 2003; Sawadogo et al., 2014; Kaindoa et al., 2019; Baeshen, 2022). While natural swarming is limited to brief dusk periods in the field, our laboratory observations here have revealed prolonged mating activities throughout the night, initiating as early as 5 PM and extending to 7 AM the following morning. The peak copulation periods were however varied slightly between *An. arabiensis* and *An. funestus*, reflecting species-specific differences. Recent field studies also support this extended mating window, with insemination rates increasing overnight, reaching ∼80–90% by early morning (Nambunga et al., 2021). These findings challenge the assumption of strictly time-bound swarming and mating behaviors, underscoring the importance of reconsidering temporal aspects of mosquito reproduction for control strategies.

The chronological age of mosquitoes was strongly associated with copulation events and mating dynamics across strains. Perhaps the most important finding of this study was that *An. funestus*, which, though poorly studied due to lack of laboratory colonies in many research groups, has significant delays in peak mating. Laboratory colonized *An. arabiensis* initiated copulation earlier post-emergence compared to *An. funestus* strains (FUTAZ and FUMOZ), reflecting species-specific differences during this period. Previous studies on *An. gambiae* and *An. stephensi* similarly reported increases in copulation success a few days post-emergence (Suleman, 1990; Oliva et al., 2011). For *An. funestus*, copulation events peaked later, likely influenced by physiological processes occurring during the pre-copulation period. Post-peak declines in mating activities were observed, possibly due to the experimental sex ratio (1 female: 2 males) and the reproductive principle that females typically mate only once (Benedict and Rafferty, 2002; Adams and Roux, 2024). These findings emphasize the importance of tailoring experimental conditions to accurately reflect natural reproductive behaviors and species-specific copulation dynamics.

Light intensity emerged as a critical factor influencing copulation dynamics. Under total darkness, observations were conducted using a flashlight for visibility, with *An. arabiensis* peaking in mating activities at 3 days post-emergence, while *An. funestus* strains FUTAZ and FUMOZ required 8–10 days. This supports previous studies that showed *An. funestus* optimal mating success is 8 days post-emergence (Maharaj et al., 2022). However, introducing a 7-watt red light significantly reduced the time to peak mating for *An. funestus*, achieving maximum copulation around 5 days. This suggests that dim red light, perhaps by simulating natural dusk conditions, likely optimizes reproductive behaviors. Studies on other *Anopheles* species, such as *An. gambiae*, similarly highlights low light as a trigger for swarming and mating, whereas excessive artificial light disrupts these behaviors (Rowland, 1989; Vielma et al., 2023; Takken et al., 2024). The association between light intensity and mating success highlights its potential for improving laboratory breeding and control interventions.

In rare observations, male-male copulation was noted in *An. funestus*. Although uncommon, such behavior likely results from sensory errors, where males misidentify fellow males as mates due to movement or wingbeat frequency (Mclver, 1980; Baeshen, 2022). This behavior may also occur because males copulate rapidly to maximize their chances of siring the next generation, leading to the clasping of other males in the swarm (Howell and Knols, 2009).

High competition, limited female availability, and altered environmental cues in laboratory settings may exacerbate these errors (Howell and Knols, 2009). While not adaptive, these instances emphasize the complexity of mating dynamics and the influence of experimental conditions on observed behaviors.

Though this study was broadly successful in achieving the set objectives, there were some minor limitations. Most importantly, the artificial laboratory conditions used in this study, including fixed light intensity and sex ratios, may limit the generalizability of the findings to wild populations. Natural variables such as fluctuating light, temperature, and humidity, as well as unexamined functions of swarming, such as navigation or predator avoidance, remain unexplored. These potential hidden functions could provide further insights into mosquito biology and vulnerabilities for vector control. For species like *An. funestus*, which are difficult to colonize, optimizing laboratory conditions, including light intensity and mating environments, is essential for colony maintenance and advancing interventions.

## Conclusion

This study sheds new light on the mating behaviors of the two malaria vectors, *An. arabiensis* and *An. funestus*, providing valuable insights into their reproductive dynamics under controlled laboratory conditions. Our findings suggest that mating in these species can extend beyond the typical dusk period, with artificial light during the night potentially having an important impact on mating behavior. The study highlights species-specific differences in reproductive timing, which could have significant implications for both wild population dynamics and laboratory-based vector control efforts. Broadly, the study showed that mating in *An. funestus* peaks much later (nearly 5 days later) than in *An. arabiensis*. Furthermore, optimizing light conditions, as demonstrated in this study, may improve the efficiency of colonizing *An. funestus*, a species notoriously difficult to breed under laboratory conditions.

Future research should focus on replicating these findings in field settings, exploring the role of additional environmental factors, and further elucidating the mechanisms behind species-specific reproductive behaviors for more targeted malaria vector control strategies.

## Acknowledgments

We would like to express our gratitude to all colleagues in the Outdoor Mosquito Control (OMC) group at Ifakara Health Institute (IHI), who provided valuable assistance throughout this work. We extend our gratitude to Steven Dominic Kipande for his dedicated technical support in monitoring the copulation experiments.

## Authors’ contributions

EEH, LLK, and FOO designed the study. EEH, ATN, DMM & JDN performed laboratory experiments. EEH, LLK, and FOO wrote and revised the manuscript. EEH performed data analysis. CAT, MJO, DJS, HSN, LLK & FOO reviewed the manuscript. All authors read and approved the final manuscript.

## Availability of data and materials

All data generated from this study will be available from the corresponding author as per request.

## Ethical approval

Ethical approval for this study was obtained from the Ifakara Health Institute Institutional Review Board with certificate number (Ref. IHI/IRB/EXT/No: 16-2024) and from the Medical Research Coordinating Committee (MRCC) at the National Institute for Medical Research-NIMR (Ref: NIMR/HQ/R.8a/Vol.IX/4112).

## Consent for publication

This manuscript has been approved for publication by the National Institute of Medical Research, Tanzania (Ref No. BD.242/437/01C/137)

